# Focal adhesion-based cell migration is differentially regulated *in vivo* versus *in vitro* by Paxillin phosphorylation

**DOI:** 10.1101/2022.03.02.482703

**Authors:** Qian Xue, Sophia R. S. Varady, Trinity Q Alaka’i Waddell, James Carrington, Minna Roh-Johnson

## Abstract

Focal adhesions are important subcellular structures that physically link the cell to the extracellular matrix (ECM), thus facilitating efficient cell migration. Although *in vitro* cell culture studies have provided a wealth of information regarding focal adhesion biology, it is critical to understand how focal adhesions are dynamically regulated in their native environment. We developed a zebrafish transplantation system in which we could efficiently visualize focal adhesion structures during single cell migration *in vivo* with high-resolution live cell imaging. By comparing focal adhesions between this *in vivo* system and the traditional *in vitro* cell culture model, we show differential regulation of a core focal adhesion protein, Paxillin. We find that a key site of phosphoregulation on Paxillin, tyrosine 118 (Y118), exhibits reduced phosphorylation in migrating cells *in vivo* in both zebrafish and mouse melanoma models, contrary to the pivotal role for this phosphorylation event in cell culture studies. Furthermore, direct modulation of this residue by site directed mutagenesis leads to opposite cell migration phenotypes *in vivo* versus *in vitro* in both migrating cancer cells and macrophages. Unexpectedly, expression of a non-phosphorylatable version of Y118-Paxillin promotes cell migration *in vivo*, despite inhibiting cell migration in the *in vitro* cell culture conditions. To further understand the mechanism of this regulation, we find that the upstream kinase, focal adhesion kinase (FAK), is downregulated in cells *in vivo*, and that cells expressing non-phosphorylatable Y118-Paxillin exhibit increased interactions between Paxillin and CRKII, an adaptor protein known to promote cell migration signaling. Collectively, our findings provide significant new insight into how focal adhesions are regulated in cells migrating in their native environment.

## Introduction

Cell migration is fundamentally required during many biological processes, including embryonic development, immune surveillance and wound healing (Horwitz and Webb, 2003; Lauffenburger and Horwitz, 1996; SenGupta et al., 2021; Trepat et al., 2012). Uncontrolled cell migration leads to diseases such as cancer metastasis and inflammation (Friedl and Wolf, 2003; Luster et al., 2005). Therefore, understanding the mechanistic basis of cell migration is critical both for fundamental aspects of biology as well as the pathology of diseases.

Focal adhesions are macromolecular structures that physically link the cell cytoskeleton to the outside extracellular matrix (ECM) during cell migration (Lauffenburger and Horwitz, 1996; Wozniak et al., 2004). The dynamic assembly and disassembly of focal adhesions at the front edge and trailing edge of the cell, respectively, are tightly coupled with actomyosin contractility to facilitate efficient cell migration (Vicente-Manzanares and Horwitz, 2011). During migration, focal adhesions are necessary for mechanical force sensation and generation, as well as serve as signaling hubs that transduce signals to the cell and to the environment (Balaban et al., 2001; Doyle et al., 2022; Eke and Cordes, 2015; Hoffman, 2014; Oakes and Gardel, 2014; Turner, 2000; Zaidel-Bar et al., 2007a). Over the past 40 years, numerous proteins have been discovered and identified as focal adhesion proteins (Burridge, 2017; Legerstee and Houtsmuller, 2021; Wozniak *et al*., 2004; Wu, 2007; Zaidel-Bar *et al*., 2007a). Furthermore, three-dimensional super-resolution microscopy techniques reveal that focal adhesions have multilaminar protein architecture, containing at least three interdependent spatial and functional protein strata, including the integrin signaling layer, force transduction layer and actin regulatory layer (Kanchanawong et al., 2010). Together, focal adhesions are well coordinated protein structures to regulate cell migration.

Most mechanistic studies of focal adhesions are based on *in vitro* cell culture assays where focal adhesions can be readily visualized at the ventral surface of cells with live cell microscopy. These mechanistic studies have identified Paxillin as a key component in focal adhesions. Paxillin acts as a scaffold in focal adhesions, transmitting extracellular signals into the intracellular space, and activating signaling cascades required for cell migration (Glenney and Zokas, 1989; Lopez-Colome et al., 2017; Turner et al., 1990). Paxillin activation is tightly regulated by the phosphorylation status of several tyrosine and serine residues along the length of the protein (Bellis et al., 1997; Burridge et al., 1992; Schaller and Schaefer, 2001). Following integrin activation, tyrosine kinases such as focal adhesion kinase (FAK) are activated and, in turn, phosphorylate Paxillin at tyrosine 118 (Y118) and tyrosine 31 (Y31) (Bellis et al., 1995; Mitra et al., 2005; Schaller and Parsons, 1995). Phosphorylated Paxillin then recruits the SH2/SH3 adaptor protein, CRKII, leading to Rac1 GTPase activation and induction of cell motility (Abassi and Vuori, 2002; Birge et al., 1993; Brugnera et al., 2002; Kiyokawa et al., 1998; Lamorte et al., 2003; Petit et al., 2000; Schaller and Parsons, 1995; Tsubouchi et al., 2002; Valles et al., 2004). Previous research also reveals that phosphorylation of Paxillin at Y118 and Y31 increases the rate of focal adhesion disassembly, thus promoting faster focal adhesion turnover and membrane protrusions in migrating cells (Zaidel-Bar et al., 2007b). Thus, tyrosine phosphorylation of Paxillin is a key event in controlling focal adhesion dynamics and overall cell migration.

A wealth of information on focal adhesions has been generated from *in vitro* 2D and 3D studies (Cukierman et al., 2001; Doyle et al., 2015; Doyle and Yamada, 2016; Fraley et al., 2010; Geiger et al., 2009; Geraldo et al., 2012; Kubow and Horwitz, 2011; Yamada et al., 2003; Yamada and Sixt, 2019). However, it is also critical to understand how focal adhesions are dynamically regulated in the *in vivo* environment in which the mechanics, signaling, and cellular interactions are intact. Focal adhesions have been shown to form and function *in vivo*, and have been best described during collective cell migration. During organ morphogenesis in *Drosophila*, focal adhesion-like structures regulate force transmission (Goodwin et al., 2016; Goodwin et al., 2017) and cell-matrix interactions (Isabella and Horne-Badovinac, 2016; Lerner et al., 2013). Similarly, in zebrafish, focal adhesion proteins are expressed in migrating sheets of cells during morphogenesis (Gunawan et al., 2019; Olson and Nechiporuk, 2021; Yamaguchi et al., 2022). During cancer progression, focal adhesion proteins can also be detected in lung metastasis in mice with fixed imaging approaches (Shibue et al., 2013). While these studies illustrate the presence of focal adhesions in collectively migrating cells, far fewer studies have examined focal adhesions *in vivo* during single cell migration *-* an important process that is highly involved in cancer metastasis and leukocyte recruitment. Live imaging studies in zebrafish show Paxillin-positive punctate structures in migrating macrophages (Barros-Becker et al., 2017), and VASP-positive punctae are found to localize to the sprouting endothelial tip cells during intersegmental vessel development (Fischer et al., 2019). However, the dynamics of focal adhesion assembly and disassembly are still entirely unclear during single cell migration *in vivo*. Furthermore, the regulation of focal adhesions is unknown in single cells migrating in a physiologically-relevant, intact environment.

A major limitation to understanding focal adhesion formation and regulation during single cell migration *in vivo* is the lack of model systems where transient subcellular focal adhesion structures form efficiently and can be readily visualized under high-resolution imaging in live animals. Here, we developed a zebrafish cancer cell transplantation system in which we can directly visualize focal adhesion structures during single cell migration *in vivo*. Taking advantage of this *in vivo* system, we aimed to determine whether new mechanisms of focal adhesion regulation could be identified in single cells migrating in their native environment as compared to the traditional *in vitro* cell culture system. Surprisingly, we found that a key focal adhesion protein, Paxillin, exhibits differential molecular dynamics in cells *in vivo* than *in vitro*. Furthermore, we found that Y118-Paxillin exhibited significantly reduced phosphorylation in migrating cells *in vivo*, despite being a key phospho-site for cell migration *in vitro*. To directly test the function of Y118-Paxillin, we performed site-directed mutagenesis studies and found that surprisingly, non-phosphorylatable Y118-Paxillin promoted cell migration *in vivo*, despite inhibiting cell migration *in vitro*. These results provide significant insight into focal adhesion regulation in cells migrating in their native environments, as well as shed light on mechanisms that affect cancer metastasis *in vivo*.

## Results

### An *in vivo* system to visualize focal adhesion dynamics

To study single cell migration *in vivo*, we took advantage of the optically transparent larval zebrafish due to the ease of cellular visualization in an intact organism. Since cell migration is a highly dynamic process that is tightly controlled by many environmental factors, we utilized a syngeneic (same species) transplantation approach in zebrafish system, allowing cells to migrate under physiological conditions. We transplanted highly migratory zebrafish melanoma cells, ZMEL cells, into the hindbrain ventricle of the larval zebrafish on 2 days post-fertilization (dpf). The larval hindbrain takes up transplanted cells readily and contains a variety of components that exist in the tumor microenvironment, such as immune cells, vasculature, ECM and other supporting cells (Roh-Johnson et al., 2017). Over the course of 1-4 days post transplantation, ZMEL cells disseminated into different larval tissue, such as the skin, neuroepithelial tissue, muscle, and tail fin (Figure 1A). In order to study focal adhesion biology during single cell migration *in vivo*, we aimed to identify an *in vivo* microenvironment where ZMEL cells use focal adhesion-based migration that is also amenable with live animal imaging. Of the regions in which ZMEL cells disseminated, we focused primarily on the skin (Figure 1B, C) as the skin is known to be enriched in ECM, a substrate that is required for focal adhesion formation. Furthermore, the skin is the relevant tissue environment for melanoma cell behavior, and it is accessible for high resolution microscopy due to its superficial location in the animal.

**Figure 1.**
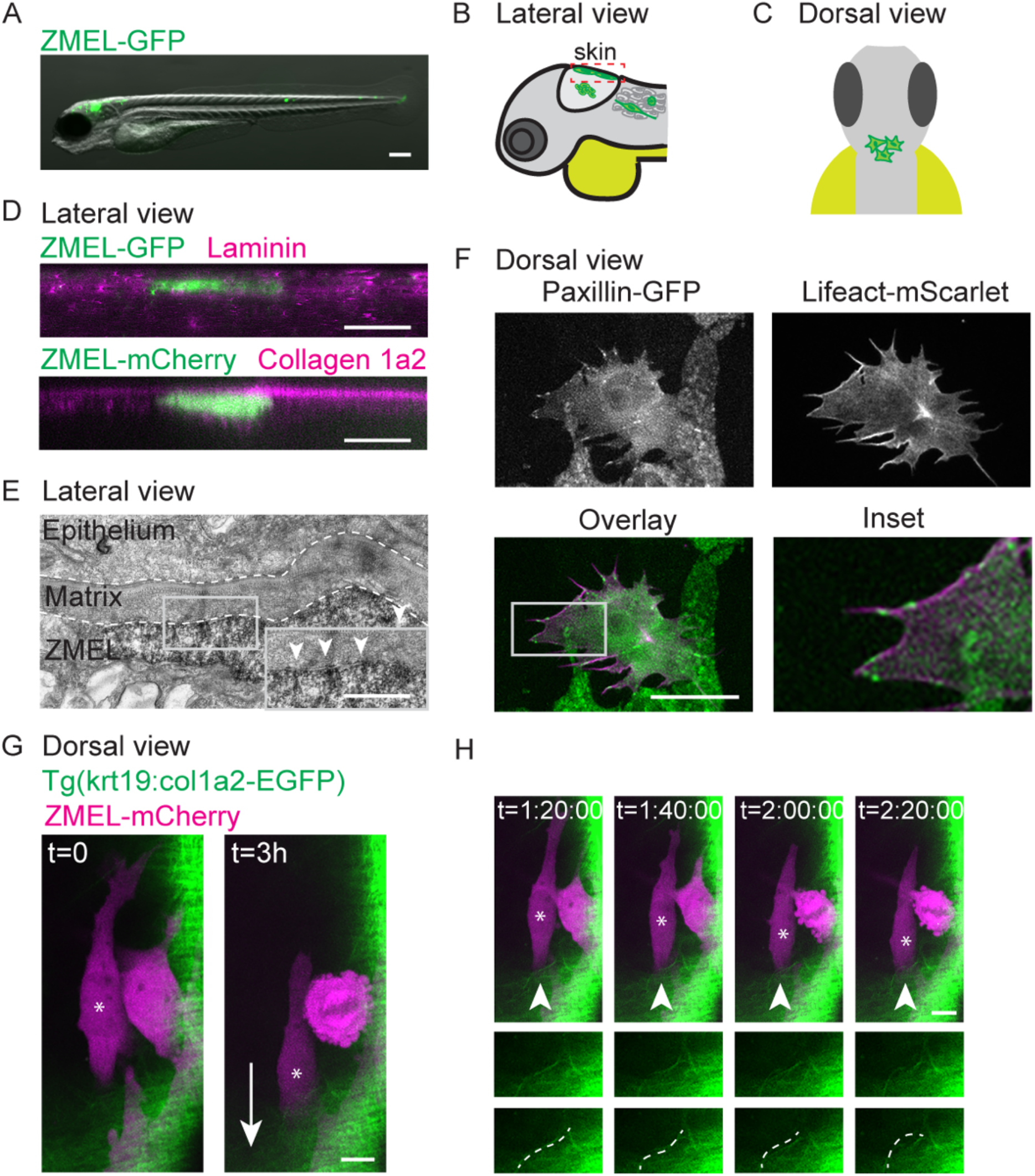
Transplanted ZMEL cells form focal adhesions structures during single cell migration *in vivo*. (A) Representative image of ZMEL-GFP whole body dissemination in a 5dpf (3 days post-transplantation) larval zebrafish. Scale bar is 100µm. (B, C) Schematic of two imaging views to visualize transplanted ZMEL cells that attach to the zebrafish skin. Lateral view (B), dashed red box indicates skin region; dorsal view (C). (D) Upper panel: Lateral view of a fixed zebrafish larva with transplanted ZMEL-GFP cells (GFP immunostaining, green) in close proximity with laminin (magenta). Lower panel: Lateral view of a live larva with transplanted ZMEL-mCherry cells (pseudocolored in green) that is proximal to collagen labeled with *Tg(krt19:col1a2-GFP)* ^*zj502*^ (pseudocolored in magenta). Scale bar is 10 µm. (E) TEM micrograph of a ZMEL cell transplanted in a larval zebrafish (3 days post-transplantation), lateral view. Dashed white lines outline the skin extracellular matrix, with a pigmented ZMEL cell underneath the matrix (labelled “ZMEL”). The inset is a magnification of the grey box revealing the ZMEL-matrix interface. Arrowheads mark the electron dense regions where ZMEL cells contact the matrix. Scale bar is 1µm. See also Figure S1 for non-ZMEL containing control larvae. (F) Live imaging of transplanted ZMEL cells co-expressing Paxillin-EGFP (Green in overlay and inset) and Lifeact-mScarlet (magenta in overlay and inset) in the zebrafish skin. Inset is magnified image of the grey box in overlay. Dorsal imaging view. Scale bar is 10µm. See also supplemental video S1. (G) Start and end frames from a timelapse video of transplanted ZMEL-mCherry cells (magenta) migrating in the zebrafish skin with collagen (green) labeled with *Tg(krt19:col1a2-GFP)*^*zj502*^. Arrow indicates the direction of migration, and the migrating ZMEL cells is marked with an asterisk. Scale bar is 10 µm. (H) Still images of timelapse video in (G). Arrowheads mark a collagen fiber (green) that is buckling as ZMEL cells (magenta, asterisk) migrate. Middle panel shows the magnified images of collagen fibers. Bottom panels highlight the buckling collagen fiber overlaid with a dashed white line. See also supplemental video S2.

We sought to determine whether ZMEL cells form focal adhesions when migrating in the skin. Thus, we first determined whether ZMEL cells migrating in the skin are in physical contact with the extracellular matrix. Using immunofluorescence assays, we found that ZMEL cells embedded in the skin are in close proximity with laminin, a protein previously reported to be enriched in skin ECM (Jessen, 2015) (Figure 1D, upper panel). Live microscopy with a collagen reporter *Tg(krt19:col1a2-GFP)*^*zj502*^ (Morris et al., 2018) revealed that ZMEL cells are proximal to the collagen-positive layer of the skin (Figure 1D, lower panel). Due to the resolution limitation of light microscopy, we further performed transmission electron microscopy (TEM) on larval zebrafish with transplanted ZMEL cells. Using non-ZMEL containing larvae as a control for overall skin tissue architecture (Figure S1A), we were able to identify ZMEL cells in the skin of transplanted larvae. TEM micrographs indicated that ZMEL cells are in direct contact with the matrices in the skin (Figure 1E). Furthermore, we visualized electron dense structures at the ZMEL-matrix interface (Figure 1E, inset), suggesting that focal adhesion-like structures form at these interfaces (Medalia and Geiger, 2010). Together, these results suggest that ZMEL cells make direct physical contact with extracellular matrix in the skin tissue of the larval zebrafish.

We next sought to determine whether ZMEL cells localize focal adhesion components to their ventral surfaces during single cell migration *in vivo*. To visualize focal adhesions *in vivo*, we utilized the zebrafish MiniCoopR system to generate zebrafish melanoma tumors expressing Paxillin (Figure S1B), a core focal adhesion protein, tagged with EGFP (Ceol et al., 2011). From these zebrafish tumors, we established a primary ZMEL cell line – ZMEL Paxillin-EGFP – using previously established approaches (Heilmann et al., 2015). When ZMEL cells are plated in the *in vitro* cell culture conditions, Paxillin-EGFP localizes to finger-like protrusions at the ventral surface of ZMEL cells, similar to what is observed in mammalian cells in culture (Turner *et al*., 1990) (Figure S1B). These Paxillin-EGFP-positive structures also assemble when ZMEL cells form protrusions, and disassemble in regions where ZMEL cells retract. These results suggest that Paxillin-EGFP localizes to focal adhesion structures in ZMEL cells in culture. To determine whether ZMEL cells form Paxillin-positive structures *in vivo*, we transplanted ZMEL Paxillin-EGFP cells into 2 dpf larval zebrafish and specifically imaged ZMEL cells in the skin. On 1 day post-transplantation, we found that Paxillin forms punctate structures on the ventral surface of ZMEL cells (the surface that makes contact with the extracellular matrix in Figure 1D) in the skin (Figure 1F). To determine whether Paxillin colocalized with another key component of focal adhesions, actin, we generated ZMEL cells co-expressing Paxillin-EGFP and Lifeact-mScarlet, and found that Paxillin-positive structures localize along actin fibers in migrating cells *in vivo* (Figure 1F; supplemental video S1). These results suggest that ZMEL cells form Paxillin-positive focal adhesion structures *in vivo*.

To determine whether ZMEL cells in the skin transduce force to the environmental ECM, we transplanted ZMEL-mCherry cells into the collagen reporter larval zebrafish. We found that as ZMEL cells migrated, a linear collagen fiber started to bend toward the migrating cell, in the opposite direction of cell migration (Figure 1G; supplemental video S2). These results suggest that ZMEL cells are actively pulling on collagen fibers as the cells migrate. Together, these results suggest that transplanted ZMEL cells form focal adhesions during single cell migration in the zebrafish skin, and we thus took advantage of this unique system of focal adhesion visualization to dissect the mechanics and regulation of focal adhesions *in vivo*.

### Paxillin exhibits distinct dynamics in migrating ZMEL cells *in vivo* as compared to migrating ZMEL cells *in vitro*

Previous studies suggest focal adhesion dynamics are affected by environmental factors and ECM (Doyle *et al*., 2015; Zhou et al., 2017). Taking advantage of this *in vivo* transplantation system, we directly compared Paxillin dynamics between *in vivo* and *in vitro* cell culture conditions. Focal adhesion dynamics are determined by the overall lifetime of a focal adhesion protein from assembly to disassembly, as well as the molecular binding dynamics which measures the molecular exchange between an adhesion and the cytosol (Stehbens and Wittmann, 2014). We first plated the primary ZMEL Paxillin-EGFP cells on *in vitro* cell culture dishes as well as transplanted the same cell line into 2 dpf larval zebrafish, and measured the molecular binding kinetics by Fluorescence Recovery After Photobleaching (FRAP). After photobleaching individual Paxillin-positive structures and monitoring the fluorescence recovery over time (Figure 2A; supplemental video S3, S4), we found that Paxillin exhibits a significantly faster molecular turnover rate in cells *in vivo* (t_1/2_=6.5±1.7s), as compared to ZMEL cells in culture (t_1/2_=15.5±4.9s) (Figure 2B). We next measured Paxillin lifetime at focal adhesions by timelapse microscopy of individual structures (Figure 2C; supplemental video S5, S6). From these measurements, we quantified overall lifetime, as well as assembly and disassembly rates as previously described (Stehbens and Wittmann, 2014) (Figure 2D). Surprisingly, we did not observe a significant difference in Paxillin lifetime between *in vivo* and *in vitro* cell culture conditions (Figure 2E); however, Paxillin exhibited significantly faster assembly rates (Figure 2F) and slower disassembly rate (Figure 2G) in cells *in vivo* as compared to the *in vitro* cell culture model. Altogether, these results indicate that although the *in vivo* environment does not affect the overall lifetime of Paxillin at focal adhesions, the assembly and disassembly rates differed between the *in vivo* and *in vitro* environments.

**Figure 2.**
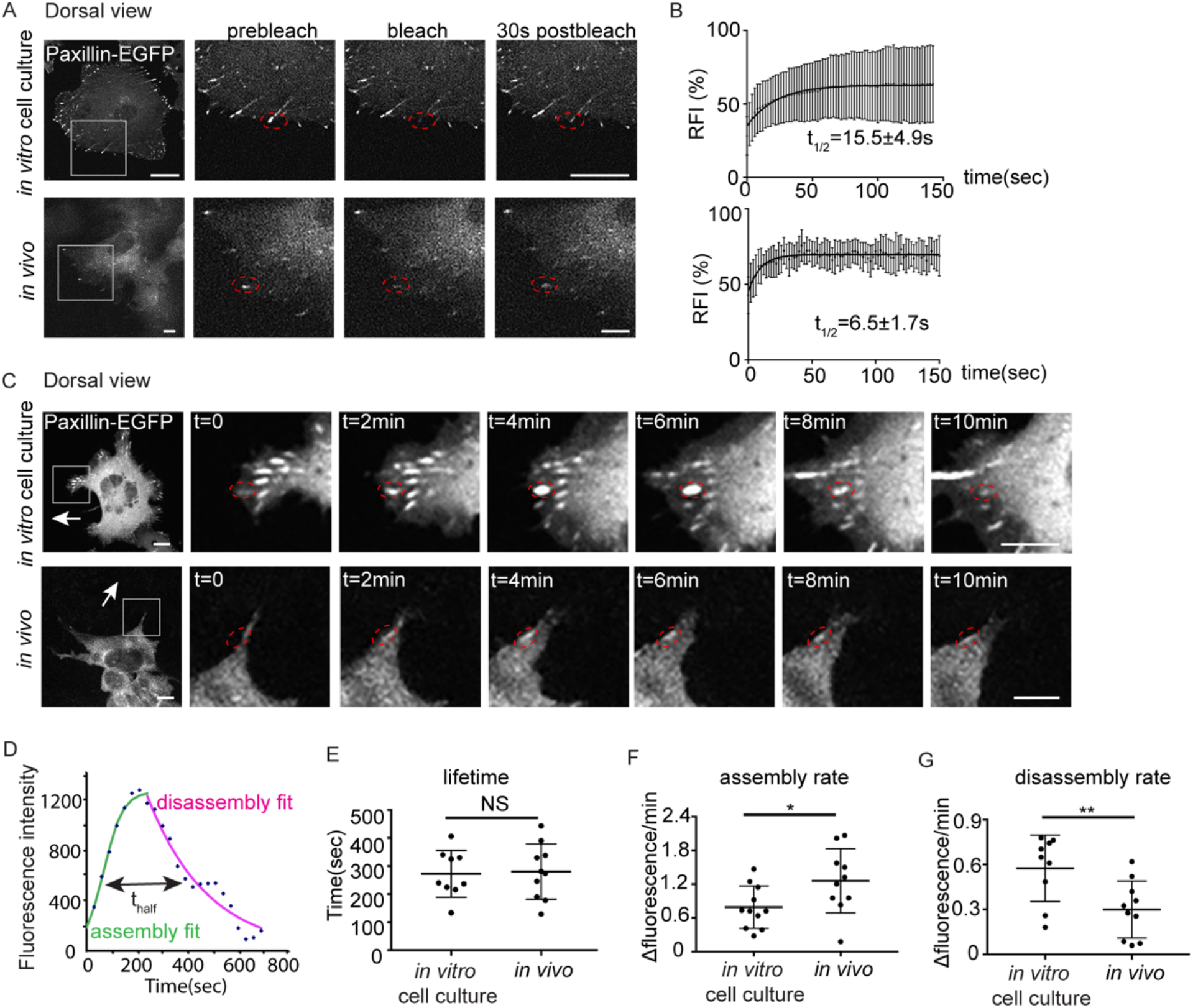
Paxillin exhibits reduced disassembly rates and increased assembly rates at focal adhesions in migrating ZMELs *in vivo*. (A) Still images of ZMEL Paxillin-EGFP FRAP experiments *in vitro* and *in vivo*. Left panel is the whole cell view and the rest of the panels are a magnification of the grey boxes prebleach, upon photobleaching, and 30 seconds after photobleaching. Red dotted circles mark Paxillin positive punctae that underwent photobleaching. See also supplemental video S3, S4. (B) Cumulative FRAP recovery curves of Paxillin-EGFP in ZMEL cells in the *in vitro* cell culture conditions and *in vivo* after photobleaching. RFI, relatively fluorescence intensity. n=36 cells for *in vitro*, n=16 cells for *in vivo* (C) Still images from timelapse videos of ZMEL Paxillin-EGFP, revealing Paxillin lifetimes at focal adhesions *in vitro* and *in vivo*. Left panel is the whole cell view and the rest of the panels are a magnification of the grey boxes. Red dotted circles mark the same Paxillin-positive punctae from assembly to disassembly. See also supplemental video S5, S6. (D) Representative graph of Paxillin lifetime curve fitting in which assembly rate, disassembly rate and lifetime (t_half_) can be calculated. (E-G) Quantification of Paxillin lifetime (E), assembly rate (F), disassembly rate (G), in the *in vitro* cell culture conditions and *in vivo* from ZMEL Paxillin-EGFP timelapse videos. n=11 cells for both *in vitro* and *in vivo*. Error bars are mean ± SD. Non-parametric unpaired t-test, *p<0.05; **p<0.01. Scale bar is 10 µm.

### Paxillin phosphorylation status is distinct in migrating cells *in vivo* versus in culture

Since we observed differential assembly and disassembly rates in Paxillin between the *in vivo* and *in vitro* cell culture conditions, we then sought to further understand the mechanisms that explain these differences. Paxillin activity has been shown to be tightly regulated by phosphorylation (Burridge *et al*., 1992), and specifically in large part through tyrosine phosphorylation by an upstream tyrosine kinase, FAK. FAK phosphorylates two tyrosine residues on Paxillin, Y31 and Y118 (Bellis *et al*., 1995; Schaller and Parsons, 1995), and phosphorylation at these sites regulates focal adhesion disassembly in cell culture models (Zaidel-Bar *et al*., 2007b). Thus, we sought to investigate whether the observed decreased rate of Paxillin disassembly *in vivo* is due to the phosphorylation status at these key tyrosine residues. We first tested the phosphorylation status of Y118-Paxillin based on its sequence conservation between zebrafish and other vertebrates (Figure 3A), as well as its previously characterized role in cell migration in other zebrafish tissue during morphogenesis (Gunawan *et al*., 2019; Olson and Nechiporuk, 2021). We examined phosphorylation of Y118-Paxillin by using a phosphospecific pY118-Paxillin antibody to stain ZMEL cells plated on *in vitro* cell culture dishes, as well as ZMEL cells transplanted in larval zebrafish. As expected, ZMEL cells plated in the *in vitro* cell culture conditions revealed clear pY118-Paxillin staining at the ventral surface of the migrating cells (Figure 3B, upper panel, white arrowheads; Figure S2A). To determine the Y118-Paxillin phosphorylation status in migrating cells *in vivo*, we performed live microscopy to first visualize migrating ZMEL cells *in vivo*, and then processed the same larvae for immunostaining to detect the pY118-Paxillin status in the same migratory cell based on the cell morphology and tissue landmarks. Surprisingly, migrating ZMEL cells *in vivo* did not have any detectable pY118-Paxillin (Figure 3B, lower panel, white arrowheads; Figure S2A). As a positive control for pY118-Paxillin immunostaining *in vivo*, we imaged the zebrafish developing heart, as previous data has indicated positive pY118-Paxillin staining in the heart (Gunawan *et al*., 2019) (Figure S2A). Furthermore, we visualized non-ZMEL cells in the skin tissue environment staining positive for pY118-Paxillin (Figure 3B, lower panel, red arrowhead), further revealing that the lack of pY118-Paxillin in ZMEL cells *in vivo* is not due to a technical issue. These findings suggest that phosphorylation of a key residue (Y118) in Paxillin that is highly enriched and essential for cell migration *in vitro*, is reduced *in vivo*.

**Figure 3.**
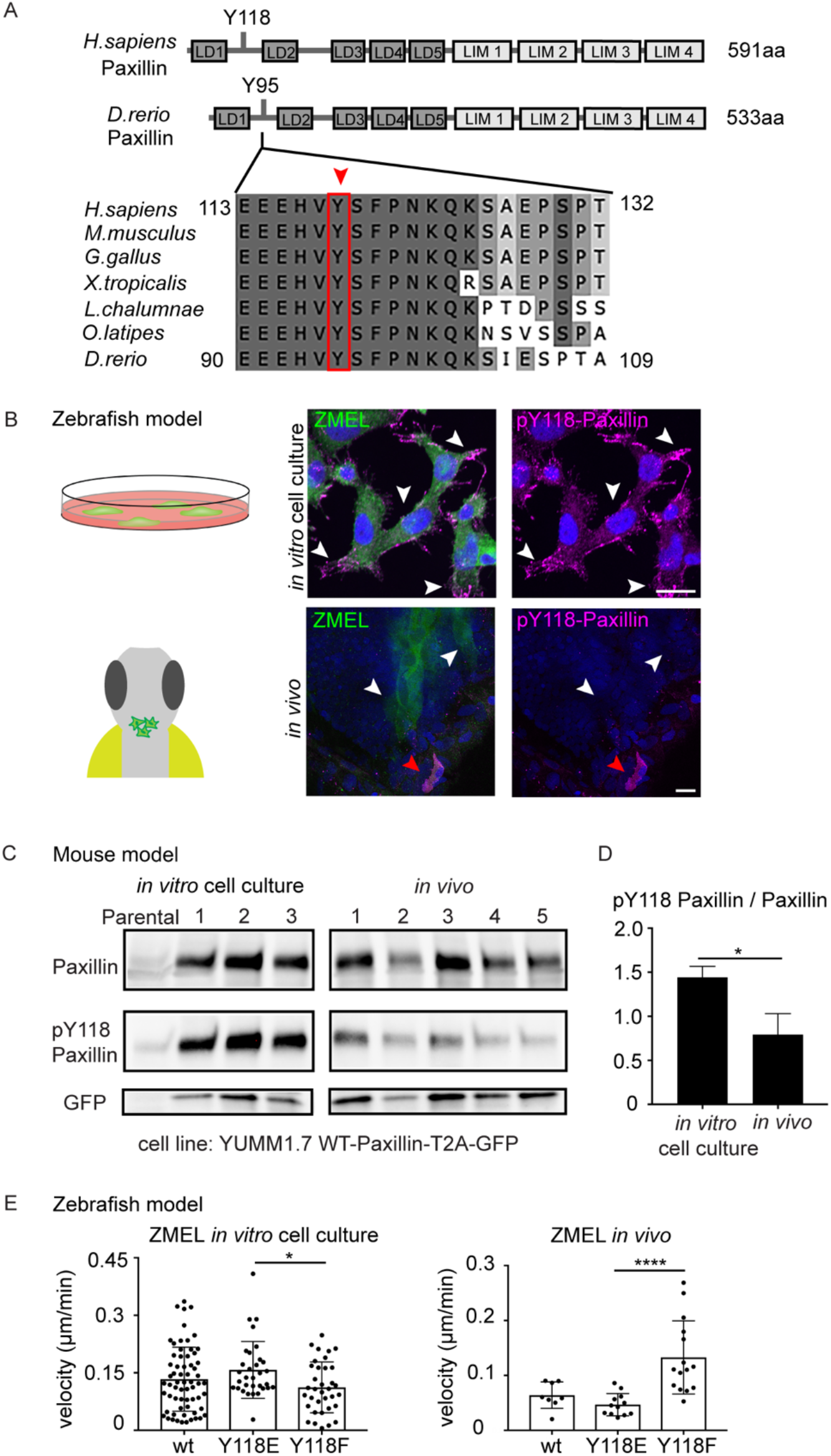
Paxillin exhibits reduced phosphorylation on Y118 in migrating cancer cells *in vivo* as compared to *in vitro* cell culture conditions in both zebrafish and mouse melanoma models. (A) Schematic of protein structures of human and zebrafish Paxillin (top) and amino acid sequence comparisons of the region encompassing Y118 between zebrafish Paxillin and vertebrate Paxillin (bottom). Red arrowhead and box indicate the conservation of Y118 Paxillin between zebrafish and other vertebrates. (B) Upper panel: pY118-Paxillin staining (magenta) of ZMEL-GFP (GFP immunostaining, green) plated on 2D *in vitro* cell culture dishes. White arrowheads mark positive pY118-Paxillin staining. Lower panel: pY118-Paxillin staining (magenta) of ZMEL-mCherry (mCherry immunostaining, pseudo-colored green, white arrowheads) in larval zebrafish (3 days post-transplantation). Red arrowhead indicates a non-ZMEL cell with positive pY118-Paxillin staining. Scale bar is 10 µm. (C) Western blot analysis of mouse melanoma YUMM1.7 cells expressing WT-Paxillin-T2A-GFP plated on the *in vitro* cell culture dishes and YUMM1.7 melanoma *in vivo* tumors. *In vitro* and *in vivo* bands are from the same blot – see entire blot in Figure S2B. GFP was used as the loading control and a control for number of YUMM1.7 cells in mouse tumors. (D) Quantification of pY118-Paxillin/total Paxillin protein ratio from (C). Non-parametric unpaired t test, *p<0.05. n=3 individual experiments. (E) Quantification of single cell migration velocity in ZMEL-mCherry cells that exogenously express GFP-tagged WT-Paxillin, Y118E-Paxillin, or Y118F-Paxillin in the *in vitro* cell culture conditions (n=64 cells for WT, n=32 cells for Y118E and n=35 cells for Y118F) and *in vivo* (n=8 cells/3 fish for WT, n=12 cells/3 fish for Y118E and n=15 cells/3 fish for Y118F). Larval zebrafish are imaged at 1 day post-transplantation. Non-parametric one-way ANOVA, *p<0.05, ****p<0.0001. Error bars are mean ± SD.

Previous studies indicate Paxillin recruitment and activation at focal adhesions are also regulated by cytoskeleton tension and integrin activation, which are processes regulated by ECM stiffness (Oakes et al., 2018; Petit *et al*., 2000). Thus, to test whether the differences in pY118-Paxillin staining is due to differential stiffness between the *in vitro* condition (glass, high stiffness) and the *in vivo* skin environment (tissue, lower stiffness), we plated ZMEL cells in a series of collagen coated dishes with stiffness ranging from 0.5kPa (brain tissue) to 50kPa (bone). Interestingly, we did not observe any lack of pY118-Paxillin staining between any of these conditions (Figure S2B), suggesting Y118 Paxillin phosphorylation in ZMEL cells is not influenced by substrate stiffness.

Due to the unexpected finding that Y118-Paxillin does not appear to be phosphorylated in migrating ZMEL cells *in vivo*, we next wanted to determine whether this phenomenon is also observable in mammalian systems. Thus, we generated melanoma tumors in mice by injecting C57BL/6J mice with YUMM1.7 cells, a highly metastatic murine melanoma cell line, expressing GFP-tagged wildtype Paxillin. We harvested tumors once they reached 1cm^3^ and performed western blot analysis to compare the Y118-Paxillin phosphorylation status of YUMM1.7 cells *in vivo* to YUMM1.7 cells in the *in vitro* cell culture conditions. Consistent with our zebrafish results, we found that YUMM1.7 cells *in vivo* consistently display a significant reduction of phosphorylation of Y118-Paxillin across all tumors we tested (Figure 3C and D; Figure S2C). These results, together with our zebrafish results, suggest that *in vivo*, migrating cells exhibit significantly reduced levels of phosphorylated Y118-Paxillin, and thus we began testing the hypothesis that lack of phosphorylation of Y118-Paxillin may facilitate cell migration *in vivo*.

### Non-phosphorylatable Y118-Paxillin promotes cell migration *in vivo*

To directly test the role of Y118-Paxillin phosphorylation during *in vivo* cell migration, we performed site directed mutagenesis to generate phosphomimetic and non-phosphorylatable versions of Y118-Paxillin. We sought to use the zebrafish *in vivo* system to test the function of these mutations due to the ability to visualize cell migration at high spatial and temporal resolution. We replaced the Y118 residue in zebrafish *paxillin* (which is residue Y95 in zebrafish *paxillin*, but will be henceforth referred to as Y118 for clarity) with a negatively charged glutamic acid (Y118E) to mimic constitutively active tyrosine phosphorylation, and replaced Y118 with phenylalanine (Y118F) to generate a non-phosphorylatable residue, as has been previously described in mammalian cells (Petit *et al*., 2000; Zaidel-Bar *et al*., 2007b). We first confirmed that these constructs localize to focal adhesion structures in ZMEL cells in the *in vitro* cell culture conditions (Figure S3). We then assessed the single cell migration velocity *in vitro* and *in vivo* of ZMEL cells expressing similar levels of GFP-WT/Y118E/Y118F-Paxillin. Under cell culture conditions, similar to what was observed in mammalian cell culture (Petit *et al*., 2000), ZMEL cells expressing the non-phosphorylatable version of Paxillin (Y118F) exhibited decreased cell migration velocities (Figure 3E, left panel). However, when ZMEL cells were transplanted *in vivo*, ZMEL cells expressing the non-phosphorylatable version of Paxillin (Y118F) exhibited faster migration velocities than both the Y118E-Paxillin-expressing cells and the wildtype-Paxillin controls (Figure 3E, right panel). Overall, these results reveal the unexpected finding that preventing phosphorylation of Y118-Paxillin has the opposite effect on cell migration *in vivo* versus *in vitro*: Non-phosphorylatable Y118-Paxillin inhibits cell migration *in vitro*, but promotes cell migration *in vivo*.

### Paxillin phosphoregulation is conserved in macrophages *in vivo*

Due to the unexpected discovery that the non-phosphorylatable Y118F mutant of Paxillin promotes ZMEL cell migration *in vivo*, despite inhibiting ZMEL cell migration in the *in vitro* cell culture conditions, we next asked whether other migrating cells *in vivo* are similarly affected by Y118-Paxillin phosphorylation status. We tested macrophages, which are highly migratory immune cells that have also been shown to use focal adhesion-based single cell migration *in vivo* (Barros-Becker *et al*., 2017). Similar to ZMEL cells, migrating macrophages in larval zebrafish do not have detectable pY118-Paxillin staining (Figure 4A). We then generated stable transgenic zebrafish strains, in which macrophages either expressed wildtype Paxillin *Tg(mpeg:WT-Paxillin-EGFP*)^*zj503*^, the phosphomimetic *Tg*(*mpeg:Y118E-Paxillin-EGFP)*^*zj504*^, or the non-phosphorylatable *Tg(mpeg: Y118F-Paxillin-EGFP*)^*zj505*^ versions of Paxillin, and crossed these strains to a macrophage reporter line *Tg(mfap4:tdTomato-CAAX)*^*xt6*^ for better macrophage visualization, to compare macrophage migration velocities during directed cell migration. We created a wound in the larval tail, and imaged macrophage migration toward the wound within 4 hours with live cell microscopy (Figure 4B). Similar to what we observed with ZMEL cells *in vivo* (Figure 3E, right panel), macrophages expressing the non-phosphorylatable Y118-Paxillin (Y118F) resulted in increased migration velocity as compared to macrophages expressing the phosphomimetic Y118E-Paxillin or wildtype-Paxillin controls (Figure 4C-E, supplemental video S7). These results demonstrate that in two distinct cell types, expressing the non-phosphorylatable version of Y118-Paxillin enhances cell migration *in vivo*.

**Figure 4.**
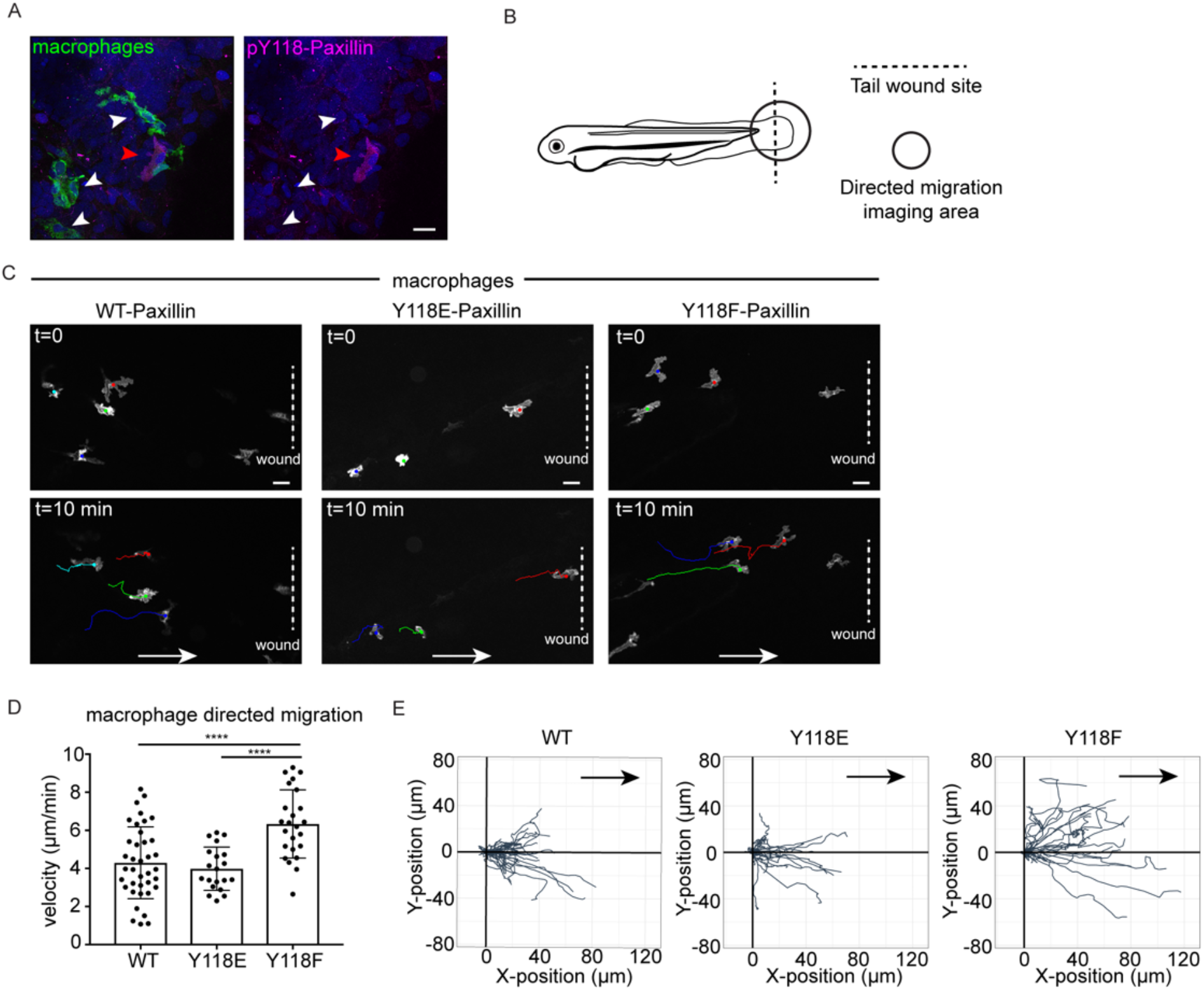
Macrophages expressing non-phosphorylatable Y118F Paxillin exhibit increased motility *in vivo*. (A) pY118 Paxillin immunostaining (magenta) of macrophages (green, white arrowheads) in *Tg(mpeg:Lifeact-GFP)*^*zj506*^ larval zebrafish. Red arrowhead marks positive pY118 Paxillin staining of a non-macrophage cell. (B) Schematic of zebrafish tail wound transection area and macrophage imaging area for directed cell migration. (C) Still images from zebrafish macrophage tracking timelapse videos in 3 dpf *Tg(mpeg:WT-Paxillin-EGFP)*^*zj503*^, *Tg(mpeg: Y118E-Paxillin-EGFP)*^*zj504*^ and *Tg(mpeg:Y118F-Paxillin-EGFP)*^*zj505*^ embryos at timepoint 0 and 10 mins. Dotted lines indicate wound sites and arrows show the direction of migration. See also supplemental video S7. Scale bar is 10 µm. (D) Quantification of macrophage migration velocities toward the wound *in vivo*. Error bars are mean ± SD. Non-parametric one-way ANOVA, ****p<0.0001. (E) Cell tracking of macrophage migration trajectories toward the wound *in vivo*, migration starting points are normalized to 0 in both x and y axes, wound sites are normalized to the positive x-axis (n=38 cells/6 fish for WT, n=20 cells/6 fish for Y118E and n=24 cells/10 fish for Y118F). Arrows show the direction of migration toward the wound.

### FAK is downregulated and CRKII exhibits increased interaction with unphosphorylated Y118-Paxillin *in vivo*

Next, we sought to understand the mechanism by which Y118-Paxillin exhibits reduced phosphorylation *in vivo*. We used mouse YUMM1.7 melanoma tumors to evaluate both upstream and downstream focal adhesion signaling. We first evaluated the well-known upstream kinase, FAK, which is known to phosphorylate Paxillin on Y118 (Figure 5A). FAK kinase activity is activated by autophosphorylation at tyrosine 397 (Y397) upon integrin-mediated cell adhesion (Bellis *et al*., 1995; Mitra *et al*., 2005; Schaller and Parsons, 1995). We first performed western blot analysis using a pY397-FAK antibody to compare FAK kinase activity of YUMM1.7 cells *in vivo* to YUMM1.7 cells that are cultured *in vitro*. Surprisingly, FAK activity (as measured by pY397FAK levels over total FAK levels) remains unchanged (Figure 5B, C; Figure S4A). However, the total FAK protein abundance is significantly decreased *in vivo* as compared to *in vitro* cell culture conditions (Figure 5B, D; Figure S4A). These results suggest that the lack of Y118-Paxillin phosphorylation *in vivo* may be due to a significant reduction in expression of the upstream kinase, FAK.

**Figure 5.**
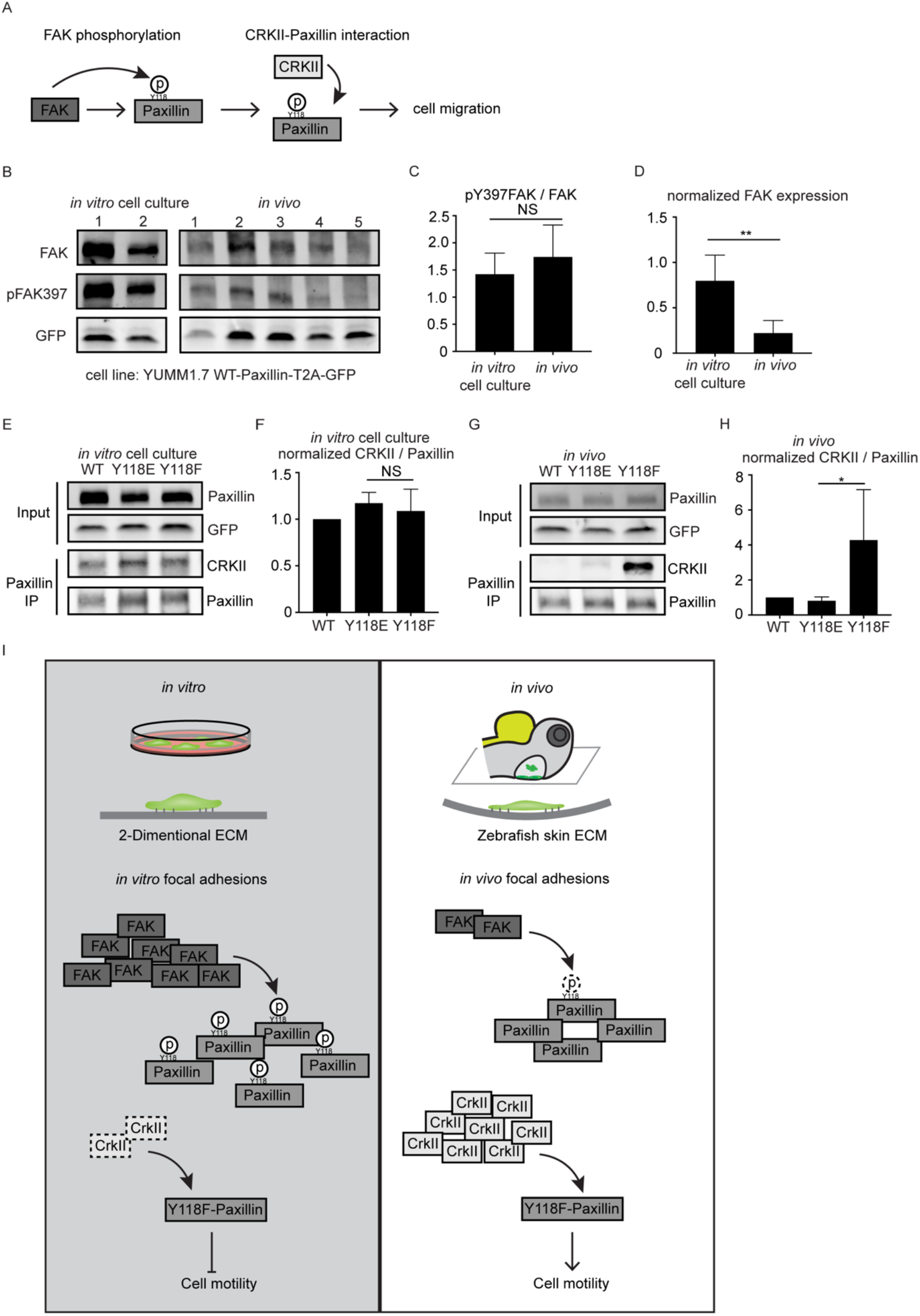
FAK is downregulated and CRKII exhibits increased interaction with unphosphorylated Y118-Paxillin *in vivo* compared to *in vitro*. (A) Schematic of *in vitro* Paxillin regulation from cell culture studies. Following integrin activation, a tyrosine kinase, FAK, phosphorylates Paxillin. Phosphorylated Paxillin then recruits the adaptor protein CRKII, activating downstream signaling pathways to induce cell migration. (B) Western blot analysis of FAK levels (FAK) and FAK activation (pFAK397) in YUMM1.7 cells expressing WT-Paxillin-T2A-GFP in culture and YUMM1.7 tumors *in vivo. In vitro* and *in vivo* bands are from the same blot. See entire blot in figure S4A. GFP was used as the loading control and as a control for the number of YUMM1.7 cells in mouse tumors. (C, D) Quantification of the pY397-FAK/total FAK ratio (C) and total normalized FAK (D) in the *in vitro* cell culture and *in vivo* conditions. Non-parametric unpaired t test, *p<0.05. n=3 individual experiments. (E-H) Co-immunoprecipitation analyses of CRKII and Paxillin in YUMM1.7 cell lines that exogenously express wildtype, Y118E and Y118F Paxillin *in vitro* (E, F) and in *in vivo* tumors (G, H). (F and H) Quantification of CRKII/Paxillin ratio from (E and G), bands from cells expressing wildtype Paxillin are normalized to 1 both *in vitro* and *in vivo*. n=3 individual experiments. (I) Working model for how Y118-Paxillin phosphorylation regulates cell migration in the *in vitro* cell culture and *in vivo* conditions.

We next aimed to dissect the underlying molecular players that promote cell migration in cells expressing the non-phosphorylatable version of Y118-Paxillin *in vivo*. We generated mouse YUMM1.7 melanoma tumors either expressing the wildtype, the phosphomimetic (Y118E), or the non-phosphorylatable (Y118F) version of Paxillin, and fused the Paxillin mutants to a self-cleavable GFP (T2A-GFP). Then, we sought to investigate interactions of Paxillin with known binding partners to see if perturbation of the Y118 residue could influence these interactions. We focused on vinculin and CRKII, which have both been shown to associate with Paxillin in response to Paxillin tyrosine phosphorylation in the *in vitro* cell culture conditions (Case et al., 2015; Pasapera et al., 2010; Petit *et al*., 2000) (Figure 5A). We first tested interactions between Paxillin and Vinculin by performing immunoprecipitations of Paxillin and assaying for Vinculin interactions. We did not observe any differences in Paxillin-Vinculin interactions in YUMM1.7 cells expressing Y118-Paxillin mutations *in vivo* (Figure S4B). However, strikingly, when we assayed for Paxillin-CRKII interactions, we found that the expression of the non-phosphorylatable Y118F-Paxillin led to significantly enhanced Paxillin-CRKII interactions *in vivo*, with no observable differences *in vitro* (Figure 5E-H). Given the critical role of CRKII in downstream cell migration (Lamorte *et al*., 2003), our results suggest that in the *in vivo* environment, non-phosphorylatable Y118-Paxillin recruits CRKII, therefore activating downstream signaling and promoting cell migration.

## Discussion

Understanding focal adhesion regulation in 3-dimensional environments and animal models is an emerging field in cell migration. In this work, we developed a zebrafish syngeneic transplantation model where focal adhesion structures can be efficiently visualized during single cell migration *in vivo*. Taking advantage of this *in vivo* approach, we performed a direct comparison of a core focal adhesion protein, Paxillin, between *in vivo* and the *in vitro* cell culture conditions (Figure 5I). Using zebrafish and mouse models, we found that Y118-Paxillin, a key residue that has been shown to be phosphorylated *in vitro*, has significantly reduced phosphorylation *in vivo*, and this lack of phosphorylation *in vivo* may be due to a significant reduction of the upstream kinase, FAK. Furthermore, preventing phosphorylation of Y118-Paxillin inhibits cell migration *in vitro*, but promotes cell migration *in vivo* in both migrating tumor cells and macrophages, and these differences correlate with the differential cellular CRKII-Paxillin interactions. Altogether, our results indicate Paxillin phosphorylation status at Y118 leads to opposite phenotypes on cell migration *in vivo* versus *in vitro* cell culture conditions.

Overexpression of Paxillin has been reported in a number of different human cancer types (Deakin et al., 2012; Mackinnon et al., 2011; Salgia et al., 1999; Sobkowicz et al., 2017; Yang et al., 2010). However, there are far fewer studies investigating Paxillin phosphorylation in cancer progression. Previous research indicates that Y118-Paxillin phosphorylation levels are inversely correlated with cancer metastasis in breast cancer patient tissue samples (Madan et al., 2006), suggesting that, similar to our findings, Y118-Paxillin phosphorylation does not correlate with an invasive phenotype. However, contrary to these findings, it has also been shown that Y118-Paxillin phosphorylation correlates with advanced human osteosarcoma metastatic stages, with highly metastatic osteosarcoma cell lines expressing high levels of pY118-Paxillin and lowly metastatic cell lines expressing low levels of pY118-Paxillin (Azuma et al., 2005). However, these experiments were performed in cell culture systems; thus, it is unclear how Y118-Paxillin is regulated *in vivo* in this system. Our melanoma work in cell culture systems is consistent with these previous *in vitro* observations, and *in vivo*, our work suggests that migrating cancer cells exhibit low levels of pY118-Paxillin, highlighting functional differences of Y118-Paxillin phosphorylation status in cancer migration *in vivo* versus *in vitro*. The dramatic increase in Paxillin/CRKII interactions in Y118F-Paxillin-expressing cells in *in vivo* mouse tumors was surprising, but it provides mechanistic support underlying the increased cell motility observed in Y118F-Paxillin expressing cells in zebrafish. CRKII has been known to directly bind Paxillin at YXXP motifs, including the phosphorylated Y118 residue of Paxillin in cultured cells (Birge *et al*., 1993; Schaller and Parsons, 1995). However, our results suggest that CRKII may bind another site on Paxillin *in vivo*, and that this interaction is enhanced in the absence of phosphorylation at Y118-Paxillin. Consistent with this hypothesis, previous data have suggested that CRKII can bind Paxillin at non-YXXP motifs (Takino et al., 2003), depending on the phosphorylation status of CRKII.

In this work, we exclusively tested the function of Y118-Paxillin in migrating cells *in vivo*. However, the phosphorylation of Y118-Paxillin is concomitant with phosphorylation at another tyrosine residue, Y31 (Schaller and Parsons, 1995). These two tyrosine residues have been largely investigated together (Petit *et al*., 2000; Tsubouchi *et al*., 2002; Zaidel-Bar *et al*., 2007b), and have been shown to play a critical role in focal adhesion turnover for efficient cell migration in cell culture (Zaidel-Bar *et al*., 2007a). To our knowledge, it is unknown whether individual tyrosine residues perform distinct functions when phosphorylated. Our work suggests that preventing phosphorylation of Y118-Paxillin alone can regulate focal adhesion dynamics during cell migration *in vivo*, and future work is required to explore the function of Y31-Paxillin.

The upstream kinase of Paxillin, FAK, is also shown to be highly expressed and activated in many types of cancers (Cance et al., 2000; Itoh et al., 2004; Madan *et al*., 2006; Miyazaki et al., 2003), and has been used as a therapeutic target for cancer treatment, including pancreatic cancer and non-small-cell lung carcinoma (Gerber et al., 2020; Parsons et al., 2008). The mechanism of FAK inhibition in cancer progression is complex, because FAK has been shown to play a critical role at focal adhesions in cell culture, and has also been shown to play signaling roles for cell survival and proliferation (Sulzmaier et al., 2014). Knowledge of FAK inhibition in regulating cancer cell migration or cancer metastasis is largely based on *in vitro* cell culture studies where FAK inhibitors alter the dynamics and formation of focal adhesion structures (Chan et al., 2009; Stutchbury et al., 2017) and downstream signals (Meng et al., 2009; Sieg et al., 2000), leading to reduced cell migration. However, it is unclear whether FAK regulates cell migration *in vivo* by the same mechanism that is described in cell culture systems, or whether focal adhesion structures are perturbed by FAK inhibitors *in vivo*, as a major limitation is the difficulty in visualizing transient and subcellular focal adhesion structures *in vivo*. In this study, we developed an animal system where focal adhesion structures can be readily visualized *in vivo*. With this system, we found that active FAK levels are reduced in cancer cells *in vivo* compared to cell culture systems, and that disrupting a key site for FAK phosphorylation on Paxillin leads to enhanced cancer cell migration *in vivo*. Together, these results suggest that FAK regulation of cancer progression *in vivo* may be mediated in large part through cell survival or proliferation signaling mechanisms, and perhaps less at focal adhesions.

Our work focuses on a single core member of focal adhesions – Paxillin. It is unknown how other focal adhesion proteins, particularly proteins from different functional layers within the focal adhesion architecture (Kanchanawong *et al*., 2010), are dynamically regulated *in vivo* and whether their regulation is distinct from the *in vitro* cell culture model. It is possible that *in vivo* focal adhesions have different multilaminar organization as compared to *in vitro* cell culture studies. Future work will be required to investigate these and other fascinating questions.

## Acknowledgments

We would like to thank all members of the Roh-Johnson lab for discussions; Rodney Stewart, Paul Martin for zebrafish reagents; members of the M. Miller and A. Welm labs for helpful discussions; the Centralized Zebrafish Animal Resources (CZAR) at the University of Utah for zebrafish husbandry and equipment; the University of Utah Cell Imaging Core for use of Leica Yokogawa CSU-W1 spinning disc confocal microscope; the University of Utah Flow Cytometry Core for cell sorting; Huntsman Cancer Institute Preclinical Research Resource for mice work; and the University of Utah Electron Microscopy Core for the help with transmission electron microscopy. This work was funded by NIH R00CA190836 (to MRJ), and 1G20OD018369-01 (to Centralized Zebrafish Animal Resource).

## Author contributions

Conceptualization, QX, MRJ; Methodology, QX, MRJ; Investigation, QX, SV, TW, JC, MRJ; Formal Analysis, QX, SV; Writing, QX, MRJ; Funding acquisition, MRJ.

## Supplemental Figures and Figure Legends

**Figure S1.**
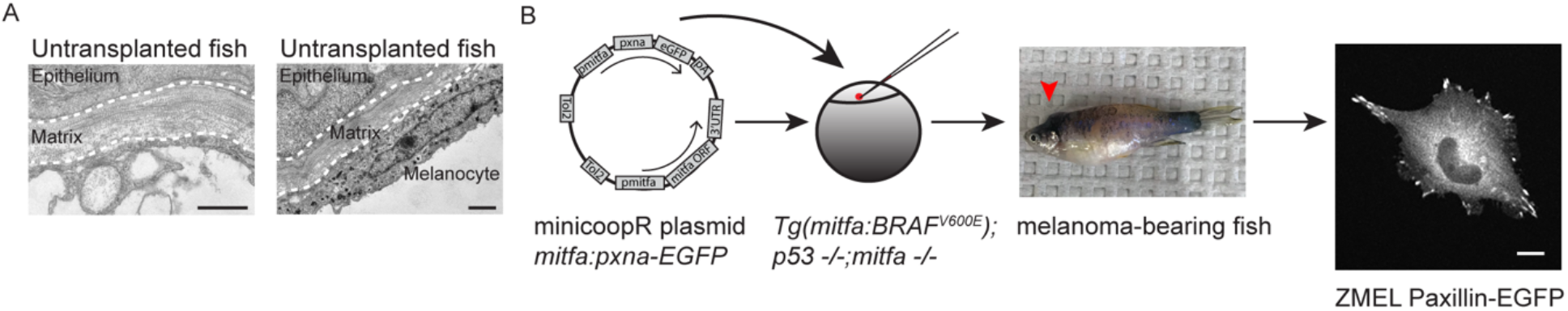
ZMEL cells form Paxillin-positive focal adhesion structures *in vitro*. (A)Two TEM micrographs of un-transplanted larvae at the skin region, 5dpf, lateral view. Dashed white lines outline the skin extracellular matrix. Left panel indicates the absence of cells underneath the matrix, and right panel indicates a pigmented melanocyte underneath the matrix. Scale bars are 1µm. (B) Schematic of the process to generate primary ZMEL Paxillin-EGFP lines. A MiniCoopR plasmid expressing GFP-tagged zebrafish paxillin-a (pxna) is injected into single-cell stage embryos of *Tg*(*mitfa*:*BRAF V600E*); *p53*(*lf*); *mitfa*(*lf*). The melanocyte rescued larvae are sorted and raised into adulthood for melanoma development (red arrowhead indicates melanoma tumor on adult zebrafish, middle panel). ZMEL Paxillin-EGFP cells are isolated from zebrafish melanoma tumors and cultured in cell culture dishes *in vitro*. Live imaging reveals Paxillin localizes to focal adhesions in *in vitro* cell culture conditions. Scale bar is 10µm.

**Figure S2.**
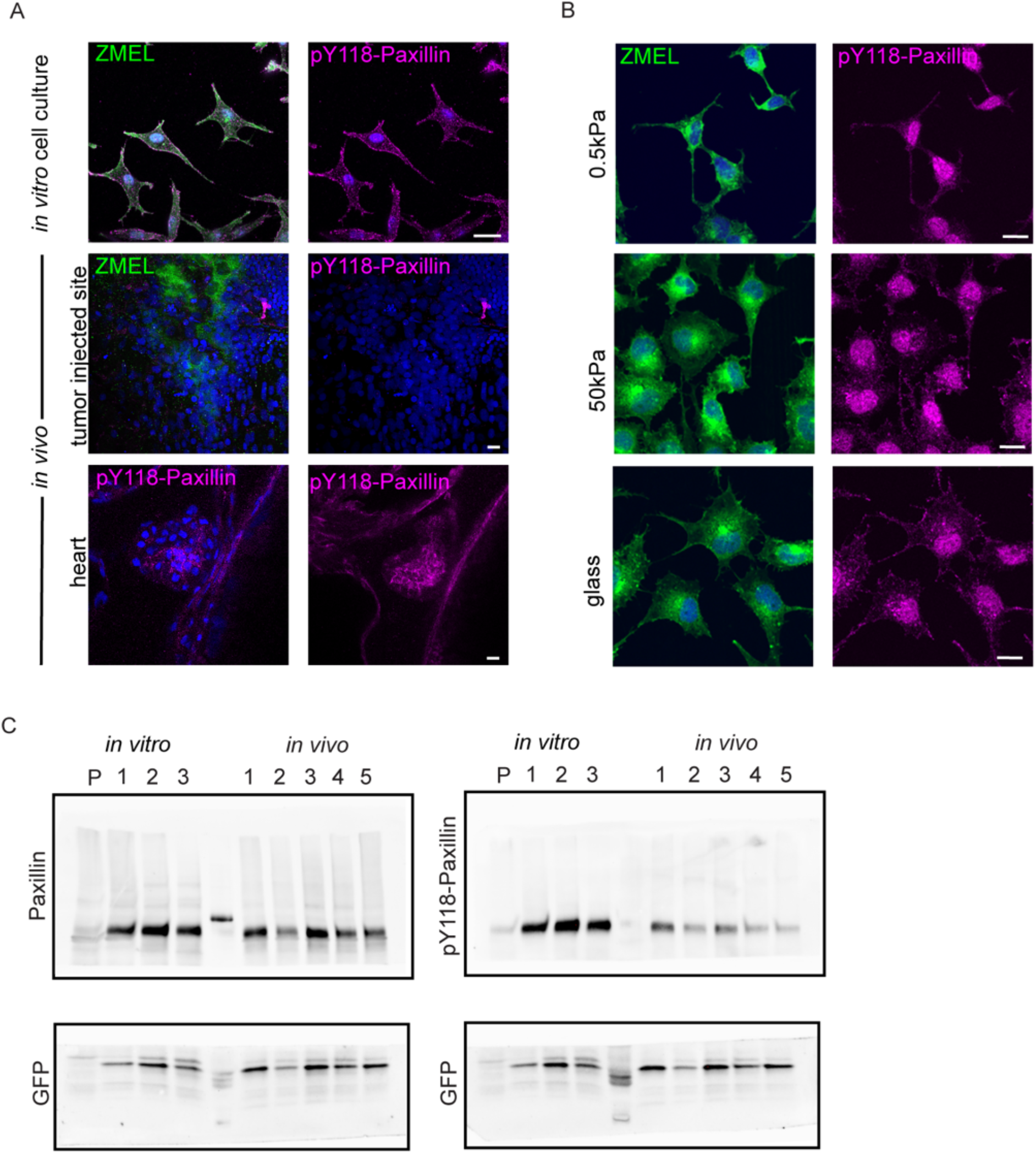
Y118-Paxillin exhibits distinct phosphorylation status in migrating cancer cells *in vivo* versus *in vitro*. (A) Upper panel: pY118-Paxillin staining (magenta) of ZMEL-GFP (GFP immunostaining, green) plated on *in vitro* cell culture dishes. Middle panel: pY118-Paxillin staining (magenta) of ZMEL-mCherry (mCherry immunostaining, pseudo-colored green) in larval zebrafish (3 days post-transplantation). Lower panel: pY118-Paxillin staining (magenta) of zebrafish developing heart (5dpf). (B) pY118-Paxillin staining of ZMEL cells expressing wildtype Paxillin plated on collagen coated dishes with different stiffness: 0.5kPa (upper panel), 50kPa (middle panel), and glass bottom dishes (lower panel). Scale bar is 10 µm. (C) Entire western blot of panels shown in Fig. 3C – YUMM1.7 cells plated in culture and YUMM1.7 melanoma tumors (*in vivo*) blotted with Paxillin and pY118-Paxillin antibodies. GFP was used as the loading control, and as a control for the number of YUMM1.7 cells in mouse tumors.

**Figure S3.**
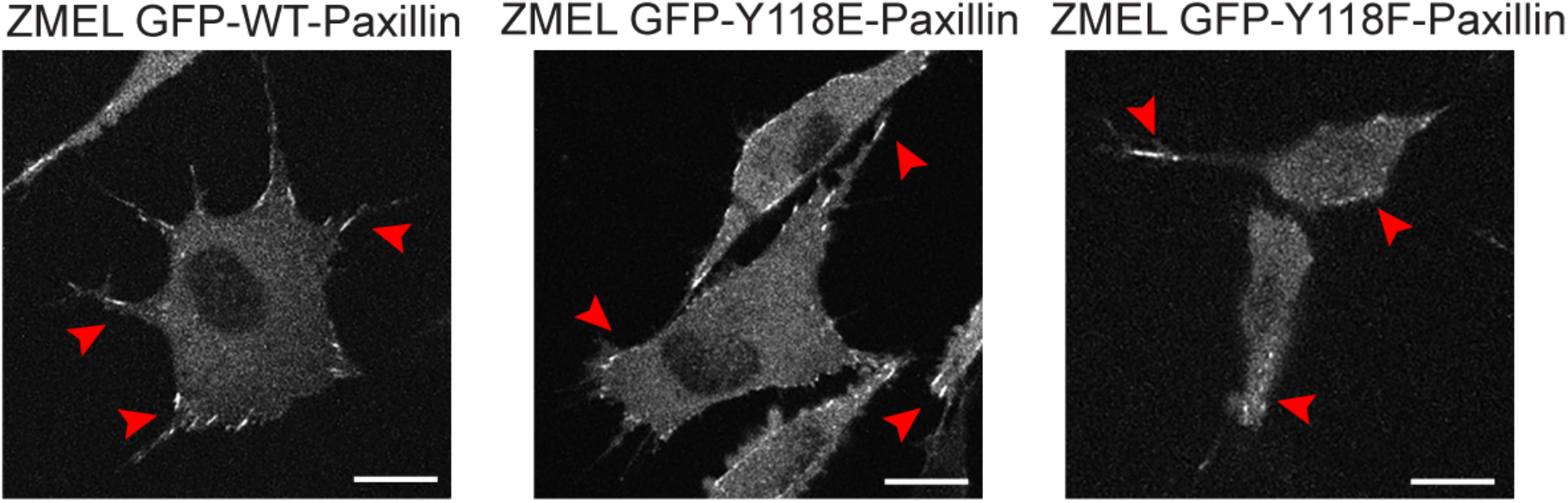
Y118-Paxillin mutants localize to focal adhesion structures in ZMEL cells in the *in vitro* cell culture conditions. Representative live images of ZMEL cells expressing GFP-WT/Y118E/Y118F-Paxillin in the *in vitro* cell culture conditions. Red arrowheads indicate Paxillin-positive focal adhesion structures. Scale bar is 10 µm.

**Figure S4.**
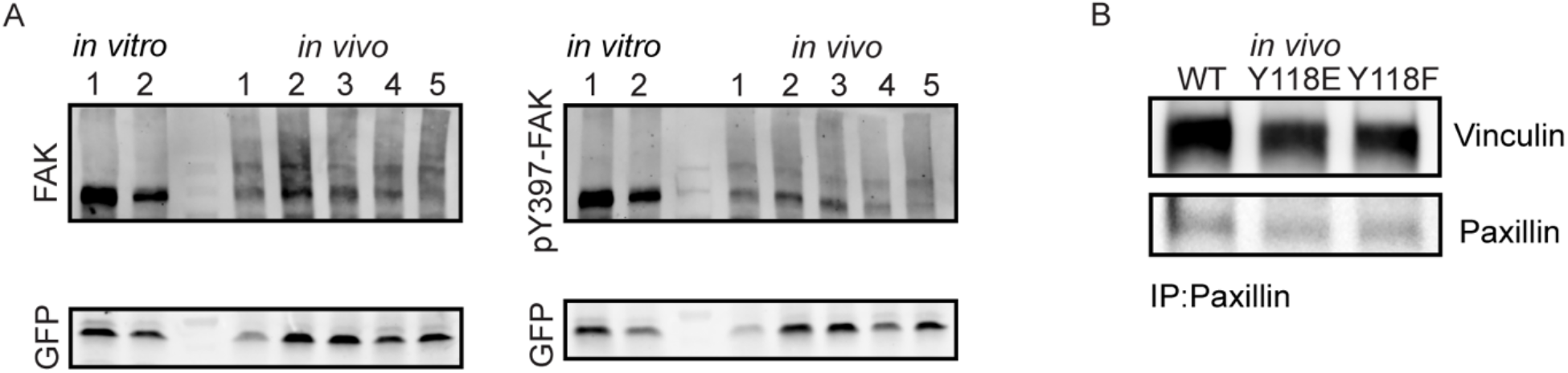
FAK is downregulated *in vivo* compared to *in vitro*. (A) Entire western blot of Fig. 5B – YUMM1.7 cells plated in culture and YUMM1.7 melanoma tumors (*in vivo*) blotted with FAK and pYFAK397 antibodies. GFP was used as the loading control, and as a control for the number of YUMM1.7 cells in mouse tumors. (B) Co-immunoprecipitation analysis of Vinculin and Paxillin, immunoprecipitating Paxillin from YUMM1.7 tumors expressing wildtype, Y118E and Y118F Paxillin and assaying for Vinculin interactions.

## Supplemental Video Legends

**Supplemental video S1: Migrating ZMEL cells in the zebrafish skin forms Paxillin-positive structures colocalizing with actin**.

Related to Figure 1. Timelapse video of a ZMEL cell co-expressing Paxillin (Paxillin-EGFP, green) and actin (Lifeact-mScarlet, magenta) migrating in the larval zebrafish skin, maximum intensity projection of Z-stacks. Images were taken every 30 seconds for 30 mins, 15 fps.

**Supplemental video S2: Migrating ZMEL cells *in vivo* transduce force to the environmental ECM**. Related to Figure 1. Timelapse video of ZMEL-mCherry cells (magenta) migrating in the skin of a zebrafish larva expressing GFP-labelled collagen *Tg(krt19:col1a2-GFP)*^*zj502*^, maximum intensity projection of Z-stacks. Arrow indicates the bending event of a collagen fiber toward the migrating cell, suggesting that the migrating cell is pulling on the collagen fiber. Images were taken every 2 minutes for 3 hours, 20 fps.

**Supplemental video S3: Paxillin FRAP in ZMEL cells *in vitro***.

Related to Figure 2. Example of a FRAP video of a ZMEL Paxillin-EGFP cell that plated in the *in vitro* cell culture conditions. A single Paxillin-positive punctum (as indicated by the red circle) is bleached, and fluorescence recovery after bleaching is recorded, single Z-plane. Images were taken every 2 seconds for 5 minutes, 7 fps.

**Supplemental video 4: Paxillin FRAP in ZMEL cells *in vivo***.

Related to Figure 2. Example of a FRAP video of a ZMEL Paxillin-EGFP cell that is transplanted *in vivo*. A single Paxillin-positive punctum (as indicated by the red circle) is bleached, and fluorescence recovery after bleaching is recorded, single Z-plane. Images were taken every 2 seconds for 5 minutes, 7 fps.

**Supplemental video S5: Paxillin lifetime in ZMEL cells *in vitro***.

Related to Figure 2. Timelapse video of a single Paxillin-positive punctum from a ZMEL Paxillin-EGFP cell that is plated in the *in vitro* cell culture conditions, maximum intensity projection Z-stacks of cell ventral surface. Images were taken every 30 seconds for 1 hour, 7 fps.

**Supplemental video S6: Paxillin lifetime in ZMEL cells *in vivo***.

Related to Figure 2. Timelapse video of a single Paxillin-positive punctum from a ZMEL Paxillin-EGFP cell that plated in the *in vitro* cell culture conditions, maximum intensity projection Z-stacks of cell ventral surface. Images were taken every 20 seconds for 1 hour, 7 fps.

**Supplemental video S7**: **Macrophages expressing non-phosphorylatable (Y118F) Paxillin exhibits increased directed cell migration toward the wound *in vivo***.

Related to Figure 4. Side-by-side timelapse videos of macrophage directed cell migration in (left) *Tg(mpeg:Paxillin-wt-eGFP)*^*zj503*^; (middle) *Tg(mpeg:Paxillin-Y118E-eGFP)*^*zj504*^; and (right) *Tg(mpeg:Paxillin-Y118F-eGFP)*^*zj505*^ zebrafish larvae, maximum intensity projection of Z-stacks. Macrophages were tracked by Manual Tracking plugin on FIJI. Images were taking every 30 seconds for 10 minutes, 7 fps.

## Materials and Methods

### Experimental models and subject details

#### Zebrafish (Danio rerio)

Zebrafish were raised in the Centralized Zebrafish Animal Resource (CZAR) at the University of Utah and experiments were approved by the Institutional Animal Care and Use Committee.

Previously described transgenic zebrafish lines used were: *Tg(mitfa:BRAFV600E);p53*−*/*−*;mitfa*−*/*−, *Tg(mfap4:tdTomato-CAAX)*^*xt6*^; and *Tg(mpeg1:lifeact-EGFP)*^*zj506*^

#### Mouse (Mus musculus)

All mouse work was performed by Preclinical Research Shared Resource at Huntsman Cancer Institute at the University of Utah, following IACUC and AAALAS guidelines. C57BL/6J mice were purchased from Jackson Laboratory (000664).

### Methods details

#### Cloning of Y118 Paxillin mutations

Zebrafish Paxillin Y118 phosphorylation mutants were constructed as previously described (Zaidel-Bar et al., 2007). Zebrafish paxillin Y95 is the ortholog of human paxillin Y118, so zebrafish paxillin 95 was replaced with glutamic acid (phosphomimetic) or phenylalanine (non-phosphorylatable). *pCS2+Paxillin-mKate* (addgene plasmid #105974) was used as the template for amplifying zebrafish paxillin-a (pxna). 1-2E/2F fragment was amplified using primer1 and primer2E/2F (Tm=71 °C) and gel purified, 3+4 fragment was amplified using primer 3 and primer4 (Tm=59 °C) and gel purified. Then the 1-2E/2F and 3+4 fragments were combined for overlap PCR using primer1 and primer4’ (Tm=55°C →70°C). *pCS2+Paxillin-mKate* was also used to amplify wildtype pxna by using primer 1 and primer4’ (Tm=70 °C). PCR products were gel purified and digested with BglII and EcoRI, and finally ligated with mEGFP-C1 (Addgene plasmid #54759) to make the final plasmids *mEGFP-WT-pxna, mEGFP-Y118E-pxna* and *mEGFP-Y118F-pxna*.

Primer1: GAAGATCTATGGACGATTTAGATGCTCTTCTCGCGG

Primer2E: CTGTTTGTTGGGGAAACTCTCGGCGTGCTCTTC

Primer2F: CTGTTTGTTGGGGAAACTGAAGGCGTGCTCTTC

Primer3: AGTTTCCCCAACAAACAG

Prmier4: GCTGAAGAGCTTGACGAAG

Primer4’: CGAATTCCTAGCTGAAGAGCTTGACGAAG

Human Paxillin Y118 phosphorylation mutants were cloned similarly as zebrafish constructs. Briefly, *pmCherry Paxillin* (Addgene plasmid #50526) was used as template. 1-2E/2F fragment was amplified using primer1-h and primer2E-h/2F-h (Tm=72 °C) and 3+4 fragment was amplified using primer3-h and primer4-h (Tm=60 °C). Then the 1-2E/2F and 3+4 fragments were combined for overlap PCR using primer1-h and primer4’-h (Tm=56°C →71°C). Wildtype human Paxillin was amplified directly by using primer1-h and primer4’-h (Tm=71 °C). PCR products were gel purified and then continued with Gibson cloning.

Primer1-h: GCCACCATGGACGACCTCGACGCCC

Primer2E-h: GCTTGTTGGGGAAGCTCTCGACGTGCTCCTC

Primer2F-h: GCTTGTTGGGGAAGCTGAAGACGTGCTCCTC

Primer3-h: AGCTTCCCCAACAAGC

Primer4-h: CTAGCAGAAGAGCTTGAGGAAG

Primer4’-h: CTAGCAGAAGAGCTTGAGGAAGCAGTTCTG

Human Paxillin fragments were then Gibson assembled with a T2A-GFP fragment into a modified pLKO.1 plasmid backbone with accessible multiple cloning sites. Briefly, Paxillin fragments were amplified from previously gel recycled products using PAX-F and PAX-R primers, T2A-GFP fragment was amplified by T2A-F and GFP-R primers using *plenti-CMV-mCherry-T2A-GFP* (Addgene plasmid #109427) as a template. Two fragments were cloned from pLKO.1 backbone, which we refer to as A1-A2 and B1-B2 fragments, by using the A1-A2 and B1-B2 primer sets listed below. Finally, Paxillin fragments were combined with T2A-GFP, A1-A2, B1-B2 fragments for Gibson assembly using NEBuilder HiFi DNA Assembly Master Mix (NEB E2621S) to generate *pLKO*.*1-WT-Paxillin-T2A-GFP*; *pLKO*.*1-Y118E-Paxillin-T2A-GFP;* and *pLKO*.*1-Y118F-Paxillin-T2A-GFP* constructs. NEB High Fidelity PCR Master Mix with HF Buffer (M0531S) was used for all PCR reactions, and sequences were verified with Sanger sequencing.

Gibson assembly primers:

A1: GTCGAGGTCGTCCATGGTAAGCTCCGGTGACGTC

PAX-F: TCACCGGAGCTTACCATGGACGACCTCGACGCCCT

PAX-R: GATCTGCACCGGGGCAGAAGAGCTTGAGGAAGCAGT

T2A-F: TCCTCAAGCTCTTCTGCCCCGGTGCAGATCTCGA

GFP-R: ACAGATATCCGTACGTTATTTATATAATTCATCCATACCGAGAG

B2: GAATTATATAAATAACGTACGGATATCTGTACAAGTAACGCCC

B1: CTTTATCCGCCTCCATCCAGTCTATTAATTGTTGCC

A2: AATTAATAGACTGGATGGAGGCGGATAAAG

### Zebrafish line generation

All constructs were generated using Gateway Cloning technology (Thermofisher), based on the manufacturer’s instructions. To generate Paxillin-expressing melanoma MiniCoopR fish, zebrafish paxillin-a (*pxna*) was cloned from the *pCS2+Paxillin-mKate* construct (Addgene plasmid #105974) using the primers listed below (pME-pxna Forward and pME-pxna Reverse). The pxna fragment was then cloned into the gateway donor vector *pDONR221(pME)* to generate *pME-pxna* by using BP clonase (Thermofisher, 11789020). Finally, *pME-pxna* was combined with *p5E-mitfa2*.*1* (addgene plasmid #81234), p3E-EGFP (gift from Rodney Stewart) and the MiniCoopR destination vector (Ceol et al., 2011) to generate the *pDest-MiniCoopR mitfa2*.*1:pxna-EGFP* plasmid using LR clonase (Thermofisher, 11791020).

Primers:

pME-pxna Forward: GGGGACAAGTTTGTACAAAAAAGCAGGCTTCGCCACCATGGACGATTTAGATGCTCTTCTC

pME-pxna Reverse: GGGGACCACTTTGTACAAGAAAGCTGGGTAGCTGAAGAGCTTGACGAAGC

Melanoma-bearing MiniCoopR fish were generated by single cell injection of *pDest-MiniCoopR mitfa2*.*1-pxna-EGFP* or *pDest-MiniCoopR mitfa2*.*1-mCherry* plasmid into embryos from *Tg(mitfa:BRAFV600E);p53*−*/*−*;mitfa*−*/*− fish with Tol2 transposase RNA as previously described (Ceol *et al*., 2011).

Collagen reporter fish were generated by single cell injection of *krt19:col1a2-GFP* plasmid (gift from Paul Martin) into wildtype zebrafish embryos.

To generate *Tg(mpeg1:WT-Paxillin-EGFP), Tg(mpeg1:Y118E-Paxillin-EGFP), Tg(mpeg1:Y118F-Paxillin-EGFP)* transgenic zebrafish lines, WT-pxna, Y118E-pxna, Y118F-pxna fragments were amplified using *mEGFP-WT-pxna, mEGFP-pxna-Y118E*, and *mEGFP-Y118F-pxna* plasmids (described above) as templates, and pME-pxna Forward/Reverse as primers. These fragments were Gateway cloned into the pME vector to generate *pME-WT-pxna, pME-Y118E-pxna* and *pME-Y118F-pxna* by using BP clonase (Thermofisher, 11789020). Then, *pME-WT/Y118E/Y118F-pxna* were combined with *p5E-mpeg1* (Roh-Johnson et al., 2017), *p3E-EGFP-pA* (gift from Rodney Steward) and *pDestpBHR4R3* (gift from Susan Brockerhoff) to generate *mpeg1: WT/Y118E/Y118F-pxna-EGFP* plasmids by LR clonase (Thermofisher, 11791020).

Transgenic embryos were all generated by single cell stage injection with each plasmid (250 ng/µL) and Tol2 transposase RNA (50 ng/µL) (Kawakami et al., 2000).

### Generation of ZMEL cell line

ZMEL cell isolation was performed as previously described (Heilmann et al., 2015). Briefly, tumors were isolated from melanoma bearing MiniCoopR fish and manually dissected in dissection medium (50% F12, 50% DMEM, 10x Pen/Strep, 0.075 mg/ml Liberase) for 30 mins at room temperature. After adding inactivating solution (50% F12, 50% DMEM, 10x Pen/Strep, 15% FBS) to the tumor suspension, tumor cells were filtered through a 40 µm strainer (Fisher Scientific, 08-771-1) 3 times and then centrifuged for 5 mins at 500 rcf and resuspended with 2ml of complete media (see details in (Heilmann *et al*., 2015)) and plated on fibronectin-coated wells of a 6-well plate. Cells were monitored for 2 weeks. Once they adhered and started to proliferate, cells were passaged to a 10 cm plate and flow cytometry was performed to sort for GFP+ cells. Primary ZMEL cells were cultured in complete media with 5% CO2 at 28.5°C. After ∼10 passages, cells began to proliferate readily, and ZMEL cells could then be cultured in standard ZMEL media (DMEM with 10% FBS and 1x glutaMAX).

### ZMEL cell transplantation into zebrafish larvae

ZMEL cell transplantations were performed as previously described (Roh-Johnson et al., 2017). Briefly, ZMEL cells were harvested and resuspended in HBSS at 10^6 cells/ml. Cells were loaded into a microinjection needle and 50 nL of the ZMEL cell suspension was transplanted into the hindbrain ventricle of anesthetized 2 dpf zebrafish larvae by using an oil-controlled microinjection rig (Eppendorf CellTram 4r Oil, #5196000030) with a Narishige arm. Injected larvae were incubated at 28.5°C. Imaging was performed 1-4 days after transplantation using a Leica Yokogawa CSU-W1 spinning disc confocal microscope in a 28.5 °C environmental chamber.

### ZMEL transfection

To exogenously express Paxillin Y118 mutants in ZMEL cells, 6 million ZMEL-mCherry cells were transfected with 30 µg *mEGFP-WT-pxna, mEGFP-Y118E-pxna* or *mEGFP-Y118F-pxna* plasmid respectively, using Neon Transfection System (Thermo Fisher; catalog #MPK10096), under 1200 V, 20 ms and 2 pulses conditions. Transfected cells were then FACS-isolated for similar expression levels of GFP.

### Zebrafish live imaging

Larval zebrafish were maintained at 28.5°C in E3 medium with 0.003% N-Phenylthiourea (Sigma Aldrich, P9629) to prevent pigmentation. For live imaging assays, 3-6 dpf larval zebrafish were anesthetized using 0.2 mg/mL Tricaine-S (Fisher Scientific, NC0872873) solution, and mounted in 1% low melting agarose (Thermofisher, #16520-050) in a 35 mm glass bottom dish (FD35-100, World Precision Instruments) for imaging. Imaging was performed using either a PL APO 40X/1.10 water immersion objective or a PL APO 63x/1.40 oil immersion objective with 1x or 2x zoom on a Leica Yokogawa CSU-W1 spinning disc confocal microscope with iXon Life 888 EMCCD camera at 28.5°C.

### Macrophage directed migration analysis

To perform directed migration assay, *Tg(mpeg1:WT-Paxillin-EGFP), Tg(mpeg1:Y118E-Paxillin-EGFP)*, or *Tg(mpeg1:Y118F-Paxillin-EGFP)* zebrafish lines were crossed with *Tg(mfap4:tdTomato-CAAX)*^*xt6*^. Tail wound transections were performed on 3 dpf larvae with a size 10 scalpel as previously described (Barros-Becker et al., 2017). Macrophage recruitment to the wound was imaged within 4 hours of wounding with standard zebrafish live imaging approaches (see above).

To analyze directed migration velocity, zebrafish macrophages were selected by thresholding the fluorescence intensity under maximum intensity Z-stacks projections, using the RFP channel, on FIJI software. The x-y coordinates of the cell centroid were tracked over time to calculate migration velocities. The final velocity was calculated by averaging the velocities from each time point of a time-lapse recording.

To plot cell migration trajectories, the x and y coordinates of macrophage centroids from the above velocity analyses were used to track macrophage migration with R (version 3.6.1) and RStudio (version 1.4.1106). Macrophage locations at timepoint 0 were normalized to the same initial coordinates, (x=0, y=0). Trajectories from fish of the same genotype were plotted in the same graph. x and y coordinates were adjusted in some fish to consistently illustrate directed migration towards the x-axis. The code for macrophage migration analysis is available on GitHub (https://github.com/rohjohnson-lab/Xue_2022).

### Focal adhesion molecular dynamic analysis

FRAP was performed on a Leica Yokogawa CSU-W1 spinning disc confocal microscope equipped with a 2D-VisiFRAP Galvo System Multi-Point FRAP/Photoactivation module. ZMEL Paxillin-EGFP cells or transplanted larvae were both imaged at 28.5°C using a Okolab stage top incubator. Single z-plane images were taken every 2 secs. 3 frames were taken prebleach, and 5 minutes of imaging were acquired postbleach. A region of interest (ROI) was drawn around individual Paxillin-positive punctae of approximately 2 µm x 2 µm for photobleaching. The FRAP laser configuration was as follows: 405nm laser line; 20 mW; 5 ms for 1 cycle.

FRAP analysis is performed as previously described (Legerstee et al., 2019). Briefly, using FIJI software, a ROI was drawn closely around bleached area of individual Paxillin+ puncta, as well as an unbleached non-cell area for background control. Mean fluorescence intensity for both bleached regions and control regions were measured. Relative fluorescence intensity (RFI) was background-subtracted and normalized to prebleached levels using: RFI=(I_t_-I_bgt_)/(I_pre_-I_bgpre_), where I_t_ is the mean fluorescence intensity of bleached area at time point t, I_bgt_ is the mean fluorescence intensity of unbleached background area at time point t. I_pre_ is the mean fluorescence intensity of the Paxillin-positive area at the prebleach timepoints; I_bgpre_ is the mean fluorescence intensity of background area at prebleach timepoints.

### Focal adhesion lifetime analysis

Focal adhesion live imaging was performed on a Leica Yokogawa CSU-W1 spinning disc confocal microscope for lifetime measurements. ZMEL Paxillin-EGFP cells or transplanted larvae were both imaged at 28.5°C using a Okolab stage top incubator. Images were taken every 20 or 30 secs for 1 hour, with 0.22 µm z-series for an approximate 5 µm z-depth.

To analyze focal adhesion lifetime, using FIJI software, z-slices comprising of 2 µm of the cell ventral surface were merged as maximum intensity projections, then a ROI was drawn closely around a focal adhesion punctum, as well as a non-cell area for background control. The mean fluorescence intensity (I_t_) of a ROI was measured at all time points from a punctum beginning to assembly until fully disassembly, and the background signal was subsequently subtracted. I_t_ was further processed with three-frame running average, and lifetime curve was generated on Excel. Curve fitting was performed as previously described (Stehbens and Wittmann, 2014). Briefly, using ‘Solver’ add-in on Excel, focal adhesion assembly curve was fit into logistic function and disassembly curve was fit into single exponential decay. Assembly and disassembly rates were calculated by the steepness of the curves and lifetime was further calculated based on the t-half of the assembly and disassembly fits.

### ZMEL cell migration velocity analysis

ZMEL cell migration was tracked by Manual Tracking plugin on FIJI software, under maximum intensity Z-stacks projection. Final velocity was calculated by averaging the velocities from each time point of a time-lapse recording.

### Immunostaining zebrafish larvae

5-7 dpf zebrafish embryos were fixed overnight at 4°C in 4% PFA. The next day, embryos were washed 3x in PBS/0.1% Tween 5 mins, and then permeabilized with 0.1% proteinase K (Thermofisher, EO0491) in PBS for 15 mins at room temperature. Embryos were then fixed in 4% PFA for 20 mins and washed with 5x with PBDT (1% BSA, 1% DMSO and 0.5% Triton X-100 in PBS) for 5 mins each. The embryos were blocked in PBDT with 10% goat serum for 2 hours before incubating in primary antibody at 4°C overnight. The embryos were then washed 6x with PBDT for 15 mins at room temperature and then incubated in secondary antibody overnight at 4°C. Embryos were dehydrated step-wise in a 25%, 50%, 75% glycerol series and were dissected and mounted for imaging. Primary antibodies used were chicken anti-GFP (Abcam, ab13970, 1:500), mouse anti-mCherry (Encor Biotechnology, MCA-1C51), rabbit anti-pY118 Paxillin (Novus Biologicals, NBP2-24459, 1:500) and Rabbit anti-Laminin (Sigma Aldrich, L9393, 1:500). Secondary antibodies used were Alexa Fluor 488 Goat anti chicken IgY(H+L) Secondary antibody (Jackson Immunoresearch, 103-545-155, 1:500), Alexa Fluor 633 Goat anti rabbit IgG(H+L) Secondary Antibody (Thermofisher, A-21070, 1:500) and DAPI (Sigma Aldrich, D9542). Imaging was performed using a 63x/1.40 oil immersion objective on a Zeiss LSM 880 (Carl Zeiss, Germany).

### Immunostaining cells in culture

One day before immunostaining, ZMEL-GFP cells and ZMEL-mCherry cells that expressing WT-Paxillin were plated on either glass bottom dishes (World Precision Instruments, FD35-100), or collagen coated dishes (Matrigen, 0.5kPa, SV3520-COL-0.5; 50kPa, SV3520-COL-50). Cells were fixed with 4% PFA and permeabilized in 0.1% Triton X-100/TBS for 5 min, then blocked with 1% BSA+1% FBS for 1 hour. Cells were incubated with primary antibodies overnight at 4°C and secondary antibodies for 1h at room temperature. Primary antibodies used were chicken anti-GFP (Abcam, ab13970, 1:500) and rabbit anti-pY118 Paxillin (Novus Biologicals, NBP2-24459, 1:500). Secondary antibodies used were Alexa Fluor 488 Goat anti chicken IgY(H+L) Secondary antibody (Jackson Immunoresearch, 103-545-155, 1:500), Alexa Fluor 633 Goat anti rabbit IgG(H+L) Secondary Antibody (Thermofisher, A-21070, 1:500), and DAPI (Sigma Aldrich, D9542). Imaging was performed using either a Zeiss LSM 880 (Carl Zeiss, Germany) with a 63x/1.40 oil immersion objective or a Leica Yokogawa CSU-W1 spinning disc confocal microscope with a Leica PL APO 63x/1.40 oil immersion objective.

### YUMM1.7 cell culture and transduction

YUMM1.7 cells were cultured in standard media (DMEM with 10% FBS and non-essential amino acids) in 5% CO2 at 37°C. To generate stably expressing YUMM1.7 cells, lentiviruses were produced by co-transfecting 293FT cells (Thermofisher, R70007) with *p*.*LKO-WT/Y118E/Y118F-Paxillin-T2A-GFP*, psPAX2 and VSV-G plasmids. Viral supernatant was collected 36 hours post-transfection and filtered through a 0.4 μm syringe filter. 50,000 parental YUMM1.7 cells were then transduced with 0.5 mL viral supernatant plus 10 μg/ml polybrene (Sigma Aldrich, TR-1003-G) on a well of a 6-well dish for 48 hours. Finally, cells were FACS-isolated for similar expression levels of GFP.

### Mouse tumor generation

YUMM1.7 cells stably expressing *WT-Paxillin-T2A-GFP, Y118E-Paxillin-T2A-GFP, Y118F-Paxiilin-T2A-GFP* were harvested and 250,000 cells were injected subcutaneously into 6-8 weeks old female C57BL/6J mice (Jackson Laboratories, 000664). Once tumors reached approximately 1cm^3, primary tumors were resected and collected for protein analysis.

### Western blot analysis

Cells in culture were lysed in RIPA lysis buffer (Thermofisher, 89900) with proteinase and phosphatase inhibitor cocktail (Sigma Aldrich, P8340, 524635) directly from cell culture plates. Tumor tissue was dissociated through ultrasonication and then lysed in the same conditions as described above. Lysates were centrifuged at 14,000 g in 4°C, and the protein concentration was measured with Pierce BCA Protein Assay Kit (Thermofisher, 23225).

15 µg protein was mixed with 4x Laemmli sample buffer (Bio-Rad, 1610747) with reducing agent and was separated by sodium dodecyl sulfate-polyacrylamide gel electrophoresis (SDS-PAGE). A 45 µm nitrocellulose membrane (Bio-Rad, 1620115) was used to transfer protein from SDS-polyacrylamide gels. Blocking buffer was either 5% BSA (in TBS/0.1% TWEEN 20) for detecting phospho-specific protein or 5% non-fat dry milk (in TBS/0.1% TWEEN 20) for non-phospho-specific protein. Primary antibodies used were Rabbit anti-Paxillin (Antibodyplus, STJ94969, 1:1000), Rabbit anti-pY118-Paxillin (Cell Signaling Technology, 9369, 1:1000), Rabbit anti-FAK (Cell Signaling Technology, 3285, 1:1000), Rabbit anti-pFAK397 (Cell Signaling Technology, 3283, 1:1000), Chicken anti-GFP (Abcam, ab13970, 1:500), Mouse anti-CrkII (BD Bioscience, 610035, 1:1000). Secondary antibodies used were Goat anti-Chicken IgY(H+L) conjugated with HRP (Thermofisher, A16054, 1:10,000), Goat anti-mouse IgG(H+L) antibody conjugated with HRP (Thermofisher, A28177, 1:10,000) and Donkey anti-rabbit IgG(H+L) antibody conjugated with HRP (GE Healthcare, NA934V, 1:10,000). Antibody signals were detected using SuperSignal West Pico Plus Chemiluminescent Substrate (Thermofisher, 34580), and were further quantified using densitometric analysis on FIJI software.

### Co-immunoprecipitation

1000 µg total protein was incubated with 5 µg Rabbit anti-Paxillin (antibodyplus, STJ94969, isotype IgG) overnight at 4°C. The next day, 1.5 mg precleared Dynabeads Protein G (10003D) were added to the protein-antibody mixture and rotated for 2 hours at 4°C, followed by 3x washes with RIPA lysis buffer. Finally, protein was eluted using 2x Laemmli sample buffer (Bio-rad, 1610747) with reducing agent. Samples were further analyzed by Western blot.

### Graphical representations and statistical analysis

All graphs were generated from Prism (v7, GraphPad), Excel (v16.43, Microsoft), R (v3.6.1, R Development Core Team 2020) and RStudio (v1.4.1106, RStudio Team 2020). Statistical analyses were performed using Prism (v7, GraphPad), R (v3.6.1, R Development Core Team 2020) and RStudio (v1.4.1106, RStudio Team 2020). We performed repeated measures ANOVA as statistical tests to take into account both the variability within the technical replicates and the biological replicates. Specific statistical tests are indicated in each figure legend.

At this time, we are not considering sex as a biological factor, as all of our mouse studies were performed in female mice, and all of our zebrafish experiments were performed in larvae that had not yet undergone sex determination.

